# Mesoscopic viscous-to-elastic transition underlies aging of biomolecular condensates

**DOI:** 10.1101/2025.11.13.688393

**Authors:** Minghao Li, Jie Lin

**Affiliations:** Center for Quantitative Biology, Peking University, Beijing, China; Peking-Tsinghua Center for Life Sciences, Peking University, Beijing, China

## Abstract

Aging of biomolecular condensates, which progressively leads to solid-like states, is closely linked to biological functions and diseases. However, a unified physical picture that captures the aging dynamics remains elusive. Here, we introduce a phenomenological model in which a condensate is a composite of viscous and elastic elements with coexisting serial and parallel connections. The model naturally yields a relaxation modulus with two relaxation times, unifying the Maxwell and Jeffreys models as special limits. When applied to aging PGL-3 condensates, our model reveals that aging is primarily driven by the conversion of viscous elements into elastic ones. In contrast, the mesoscopic structure topology of the condensates remains approximately invariant. Our stochastic theory of element dynamics also predicts that the condensate viscosity either increases exponentially, in agreement with experiments, or as a power law of the waiting time, governed by a single parameter that depends on temperature and the molecular interaction strength.

## Introduction

Biomolecular condensates formed through liquid-liquid phase separation are essential for cellular organizations and functions [1–11]. Theoretical and experimental studies have provided profound insights into the molecular interactions governing the formation of condensates [12–17]. Meanwhile, biomolecular condensates often undergo aging [18–26], progressively hardening into solid-like states, or even pathological aggregates associated with neurodegenerative diseases [27–30]. Therefore, understanding the physical principles governing the transition in rheological properties during the aging of biomolecular condensates is extremely important. Recent experiments have characterized biomolecular condensates as viscoelastic materials, fitting the dynamic modulus data using the Maxwell model [31] or the Jeffreys model [32]. Nevertheless, it is still unclear whether a theoretical framework can unify these experimental observations and quantify the mechanical states of biomolecular condensates during aging.

Previous studies have proposed various microscopic mechanisms for the aging of biomolecular condensates from different perspectives. For instance, molecular dynamics simulations employing all-atom or coarse-grained approaches have been used to compute intermolecular interactions and extract aging dynamics [14, 15, 33–37]. However, a unified physical picture of condensate aging remains lacking. On the other hand, phenomenological theories have been proposed in which the condensate is composed of mesoscopic elements, and aging is attributed to the change in the energy-barrier distribution of the traps these elements inhabit. These theories predict temporal scaling for the viscosity, *η* ∼ *t*^1−*α*^ where *α* = 0 [38] or 0 *< α <* 1 [39] where *t* is the waiting time since droplet formation. However, experiments showed that viscosity increases approximately exponentially with the waiting time [31]. Furthermore, these theories predict a single relaxation time for the viscoelastic response of biomolecular condensates [38, 39]. However, experimental data on the dynamic modulus appear to be better described by more complex models with two relaxation times [31, 40].

In this work, we introduce a viscoelastic model of biomolecular condensates, offering a unified phenomenological description of the rheological properties of biomolecular condensates. Our model conceptualizes a condensate as a composite material composed of viscous and elastic elements. Importantly, we assume that a finite fraction of these elements are connected in parallel, while the rest are connected in serial, which reflects the mesoscopic structure topology of the condensate. Intriguingly, our model naturally predicts a relaxation modulus with two relaxation times. Our theory reconciles the previous Maxwell and Jeffreys descriptions as limiting cases of our model [31, 32], and also provides a superior fit to experimental data. We also provide a kinetic model to describe the time dependence of the fraction of viscous elements (*P*). Motivated by experiments and simulations that show the aging of protein condensates is driven by the slow accumulation of strong interactions, notably the formation of *β*-sheets [15, 24, 25, 34, 37], we introduce a deep energy barrier representing these stable structures alongside a population of exponentially distributed barriers representing transient weak interactions. Notably, this minimal model successfully explains the exponential increase in viscosity over time.

In the following, we first introduce our viscoelastic model and demonstrate that the condensate’s viscoelasticity is characterized by a modified Maxwell model with two relaxation times that depend on the fractions of viscous and serial elements. We then apply our model to data of aging PGL-3 condensates [31]. Remarkably, we find that the aging of PGL-3 condensate is not driven by a change in the mesoscopic structure topology (the fraction of serial connections, *f*, remains approximately constant), but rather by the progressive conversion of viscous elements into elastic ones (the fraction of viscous elements, *P*, decays approximately exponentially). This conversion elucidates the mesoscopic origin of the exponential increase in viscosity, as our model predicts a simple relationship, *η* ∝ 1*/*(*f P*).

We then introduce a kinetic theory for the aging behaviors of the fraction of viscous elements. The theory predicts two distinct aging modes: the viscosity may increase exponentially with time or grow as a power-law function of time *η* ∼ *t*^1+*α*^ with 0 *< α <* 1 in the long-time limit. Importantly, the transition in the viscosity scaling law can be triggered by varying temperature. Our framework thus provides a bridge from mesoscopic state transitions to the macroscopic material fate of biomolecular condensates, offering insights into the physical principles underlying their aging.

### Viscoelastic model

We divide the biomolecular condensates into mesoscopic elements, whose sizes are sufficiently small so that one condensate is composed of numerous elements, yet large enough so that their mechanical response can be quantified by strain and stress. We assume that each element is either a viscous element with constant viscosity *η*_0_ or an elastic element with constant shear modulus *G*_0_. The fraction of viscous elements is *P*. In the soft glassy rheology model [41, 42], multiple Maxwell-fluid elements are connected in parallel, sharing the same strain and contributing together to the stress. In the model by Takaki et al. [39], elements are connected in series, sharing the same stress and contributing together to the strain.

Here, we introduce a viscoelastic model in which serial and parallel connections coexist, with the fraction of elements in the serial section denoted by *f* (Figure 1). As the condensate ages, the fraction of viscous elements *P* and the fraction of elements in the serial section *f* may change. The serial and parallel sections satisfy the following constitutive relationship:

**FIG. 1.**
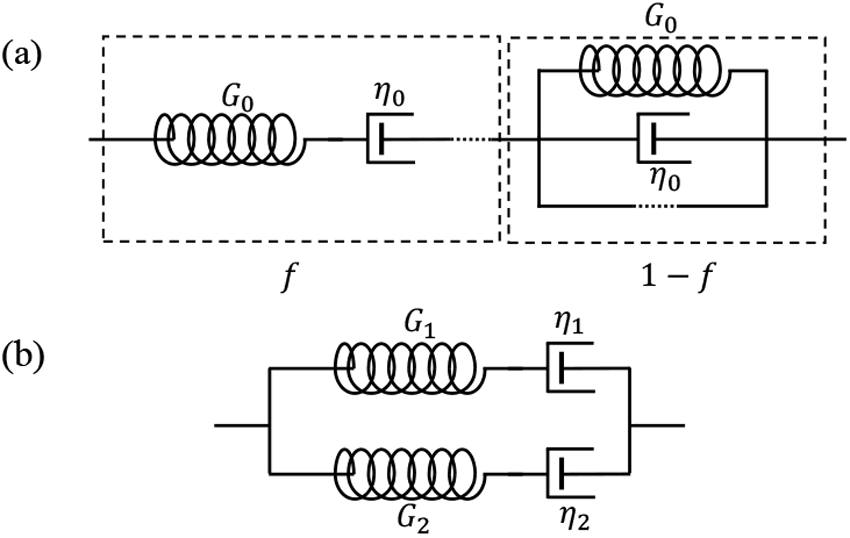
(a) A phenomenological model of biomolecular condensate rheology. The model consists of elastic and viscous elements, represented here by springs and dashpots connected in series or in parallel. We use the ellipses to indicate unshown elements. A fraction *P* of the elements are viscous elements, and a fraction *f* of the elements are in the serial section. (b) The viscoelastic response of the model in (a) is equivalent to two Maxwell fluids with different shear moduli and viscosities connected in parallel.

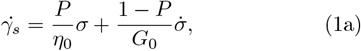

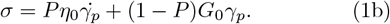

Here, *σ* is the stress, and *γ*_*s*_ and *γ*_*p*_ are the strains of the serial and parallel sections, respectively. The total strain of the whole condensate, *γ* = *fγ*_*s*_ +(1 − *f*)*γ*_*p*_. We remark that the system size should not appear in Eqs. (1a, 1b) since strain and stress are intensive variables.

The relaxation modulus of the above viscoelastic model (i.e., the stress relaxation given a step unit strain) is precisely a double-exponential decay function with a short relaxation time *τ*_1_ and a long relaxation time *τ*_2_ ( Figure 2a and see detailed derivation in Supplemental Material):

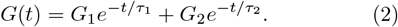

**FIG. 2.**
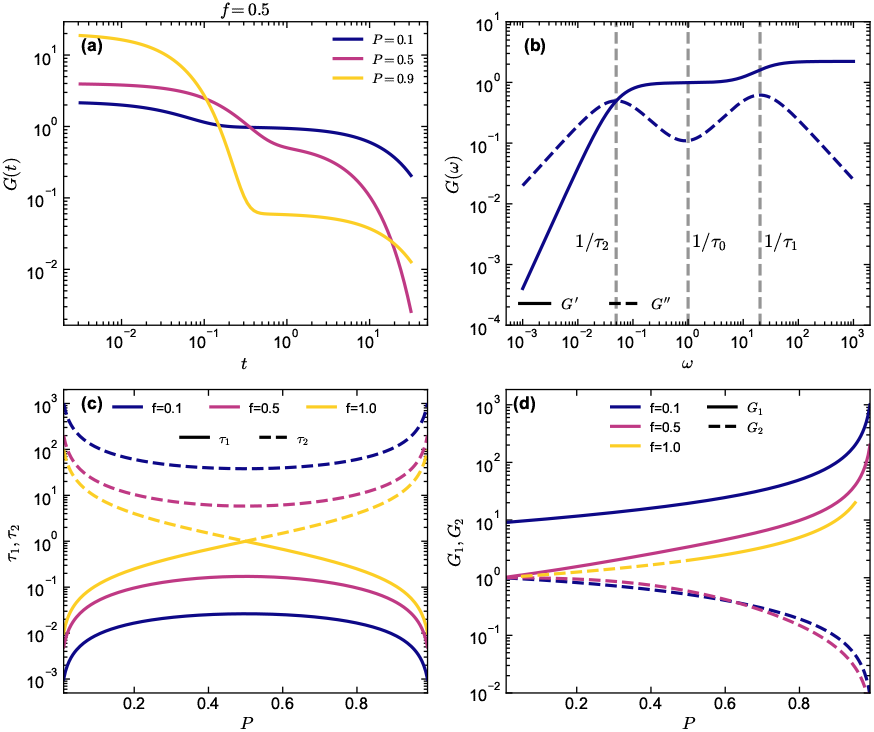
(a) The relaxation modulus *G*(*t*) vs. *t* for different values of the viscous fraction *P* at a fixed serial fraction *f* = 0.5. (b) The storage (*G*^*′*^, solid lines) and loss (*G*^*′′*^, dashed lines) moduli vs. frequency *ω* for *f* = 0.5, *P* = 0.1. We highlight the frequencies corresponding to the slow mode 1*/τ*_2_, the fast mode 1*/τ*_1_, and the mesoscopic time scale 1*/τ*_0_. In the limit *ω* → 0, *G*^*′*^ ∝ *ω*^2^ and *G*^*′′*^ ∝ *ω*. → ∞ In the limit *ω, G*^*′*^ 1*/*[*f* (1 *P*)] and *G*^*′′*^ ∝ *ω*^−1^. (c) The relaxation times *τ*_1_ and *τ*_2_ as functions of *P* for different values of *f*. For *f* = 1, the curves of *τ*_1_ and *τ*_2_ intersect at *P* = 0.5. (d) The amplitudes *G*_1_ and *G*_2_ as functions of *P* for different *f*. In this figure, the modulus unit is *G*_0_ and the time unit is *η*_0_*/G*_0_.

Here, the two relaxation times and their corresponding amplitudes are given by

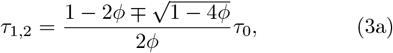

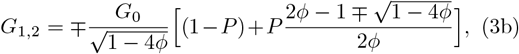

where *τ*_0_ = *η*_0_*/G*_0_ and *ϕ* = *f P* (1 − *P*) (Figure 2c, d). Notably, the relaxation times exhibit the symmetry 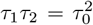 (Figure 2b). In particular, *τ*_1_ ∝ *f P* and *τ*_2_ ∝ 1*/*(*f P*) as *P* → 0. Given the relaxation modulus, it is straightforward to compute the zero-shear viscosity and the instantaneous modulus:

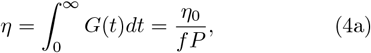

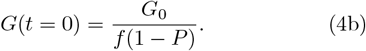

The condensate becomes a solid when *f* = 0, that is, all elements are connected in parallel; in this case, the plateau modulus *G*_∞_ = (1 − *P*)*G*_0_. To summarize, the fraction of viscous elements, *P*, and the fraction of serial elements, *f*, uniquely determine the viscoelastic behaviors of condensates. In the following, we apply our theory to fit the dynamic moduli of protein condensates at different waiting times after condensate formation.

### Analysis of experimental data

Our viscoelastic model predicts that the dynamic modulus of the condensate is equivalent to two Maxwell fluids connected in parallel

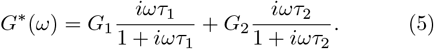

Previous works have suggested that the dynamics moduli of biomolecular condensates can be described by the Maxwell model [31] and the Jeffreys model [32]. Our model unifies the two different hypotheses. There are two possible scenarios that Eq. (5) reduces to a Maxwell model: (1) *τ*_1_ ≈ *τ*_2_, i.e., the two relaxation times are close; (2) the loss modulus *G*^*′′*^(*ω*) is still decreasing due to the slow mode even at the largest frequency *ω*_max_ the experiment can probe, i.e., *G*_2_*/*(*w*_max_*τ*_2_) *> G*_1_*ω*_max_*τ*_1_, which means that 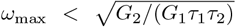. For Eq. (5) to resemble Jeffreys model, we require that *G*^*′′*^(*ω*) starts to increase due to the fast mode at high frequ<u>encies but d</u>oes not reach the second peak yet, i.e.,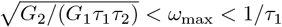 (Figure 2b).

We use Eq.(5), together with Eqs. (3a, 3b) to fit the dynamic moduli data of PGL-3 protein measured by active microrheology [31]. The fitting parameters are *P, f, G*_0_, and *η*_0_. The fitting results suggest that our model can well fit the aging PGL-3 condensate with two different relaxation times (Figure 3a-f). Interestingly, we observe a relatively constant fraction of serial elements *f* (*t*) (Figure 3i), which suggests that the condensate’s mesoscopic structure topology does not change significantly during aging. Meanwhile, we observe an approximately exponential decay of the fraction of viscos elements *P* (*t*) (Figure 3j).

**FIG. 3.**
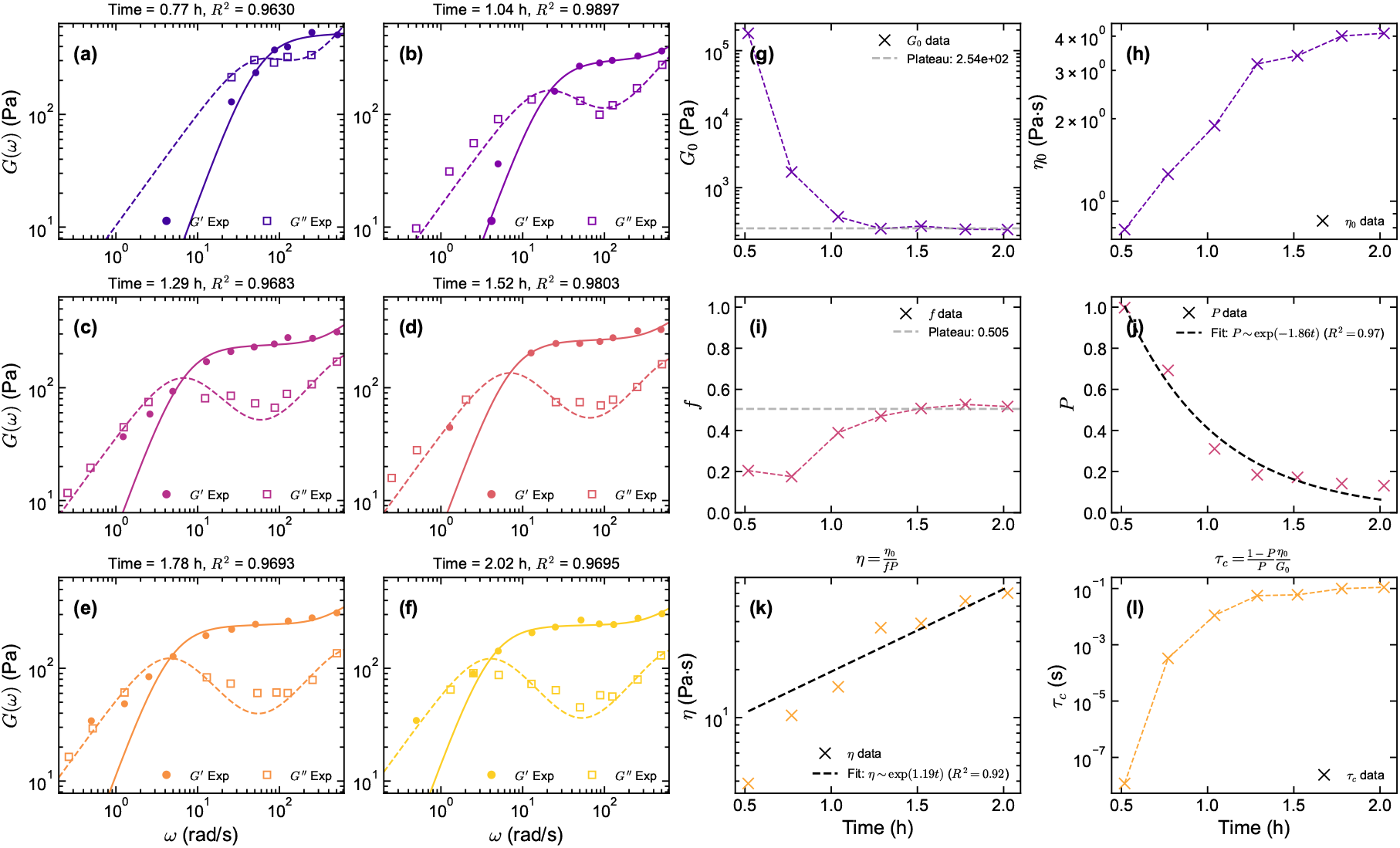
(a-f) Fitting of active microrheology data for PGL-3 condensates. The solid (*G*^*′*^) and dashed (*G*^*′′*^) lines are fits to our model predictions, Eq. (5). (g-j) The fitting parameters *G*_0_, *η*_0_, *P*, and *f* in (a-f) at different waiting times. *G*_0_ and *f* reached plateaus at 254 Pa and 0.505 (grad dashed lines), respectively. *P* vs. *t* can be well-fitted by an exponential function (dashed line). (k-l) The time dependence of *η* and *τ*_*c*_ calculated from the parameters in (g-j). The viscosity increases approximately exponentially in time.

We calculate the viscosity *η* of the condensate by Eq. (4a), and the relaxation time as *τ*_*c*_ = *η/G*(*t* = 0) (Figure 3k, l). Notably, the inferred viscosity *η* = *η*_0_*/f P* also increases approximately exponentially since its value is dominated by the fraction of viscous elements *P* while the change of *η*_0_ is milder (Figure 3h). Notably, *G*_0_ of the elastic elements undergoes a fast decay in the early stage and reaches a plateau in the later stage (Figure 3g). Our analysis suggests that aging of PGL-3 condensates primarily stems from changes in *P* and *G*_0_. In the following, we explain the mechanism of the exponential decay of *P* (*t*).

### Aging dynamics of condensates

We introduce a stochastic model to describe the time dependence of the fraction of viscous elements *P* (*t*). We introduce *p*_*e*_(*E, t*) for the distribution of elastic elements as a function of their energy barriers, which satisfies 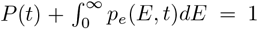. The change of the state of each element is stochastic, governed by the following Master processes:

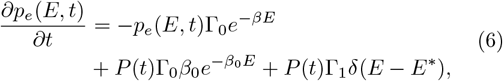

where *β* ≡ *k*_*B*_*T*, with *k*_*B*_ the Boltzmann constant and *T* the temperature. Here, the elastic state is further divided into two substates. The weakly elastic substate, characterized by an exponential distribution of energy barriers 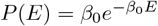, represents transient and weak interactions. In contrast, the strongly elastic substate, described by a delta function *P* (*E*) = δ(*E* − *E*^∗^) with *E*^∗^ ≫ *k*_*B*_*T*, represents the deep energy trap associated with the slow accumulation of stable structures, such as *β*-sheets, which drive the liquid-to-solid transition [15, 24, 33]. For both substates, the transition rate from them to the viscous state follows the Arrhenius law, Γ = Γ_0_*e*^−*βE*^. We introduce the parameter *α* ≡ *β*_0_*/β* and summarize the temporal scaling behavior of the viscous fraction in the following (see detailed derivation in Supplemental Material).

For *α >* 1, starting from the initial condition *P* (*t*) = 1, i.e., a fresh condensate, *P* (*t*) first enters a plateau regime *P* (*t*) = (*α* − 1)*/*(2*α* − 1) and then decays exponentially as

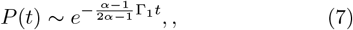

in agreement with the experimental measurements. The transition between the plateau and the exponential decay occurs at *t*_*c*_ ∼ 1*/*Γ_1_ (Figure 4a, b). For *α <* 1, *P* (*t*) decays as a two-stage power-law

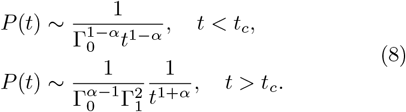

**FIG. 4.**
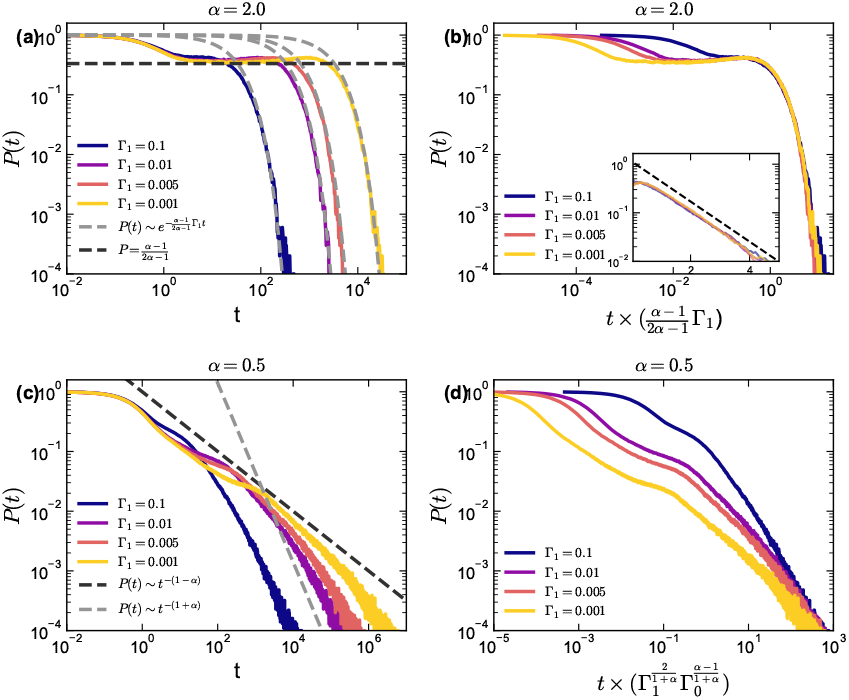
Aging dynamics of the fraction of viscous elements. (a) We compare the simulation results (solid lines) with the theoretical predictions (dashed lines). *P* (*t*) first reaches a plateau and then decays exponentially for *α* = 2. (b) The long-time behaviors of the *P* (*t*) vs. *t* curves collapse on a master curve after rescaling the time as 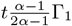. (c) The same as (a) but for *α* = 0.5 in which *P* (*t*) transitions between two power-law decay regimes. (d) The same as (c) but with a rescaling factor. In this figure, we take *N* = 100000 elements in the simulations. We set *E*^∗^ = 30, the energy unit as *k*_*B*_*T*, and the time unit as 1*/*Γ_0_.

Here, the crossover between the two power-law regimes occurs at 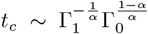 (Figure 4c, d). To confirm our predictions, we simulate the stochastic model using the Gillispie algorithm and find perfect agreement (Figure 4). We remark that a power-law function can also fit the experimental data of *P* (*t*) (Figure S1), and our model predicts that the exponent must be larger than 1, in agreement with the inferred value, 1 + *α* ≈ 1.56.

## Discussion

In this work, we introduce a phenomenological viscoelastic model that provides a unified framework for understanding the complex rheology and aging dynamics of biomolecular condensates. We demonstrate that a simple model incorporating both serial and parallel arrangements of viscous and elastic elements yields a double-exponential relaxation modulus, with the Maxwell and Jeffreys models as limiting cases. By fitting our model predictions to the data of PGL-3 condensates [31], we decompose the aging process into the time evolution of four physical parameters: the fraction of viscous elements (*P*), the fraction of serial connections (*f*), and the intrinsic properties of the elements (*G*_0_, *η*_0_).

Our analysis reveals an important insight: the mesoscopic structure topology of the PGL-3 condensate network, characterized by a nearly constant *f* ≈ 0.5, appears to be established early and remains stable throughout aging (Figure 3i). In contrast, aging is predominantly driven by the conversion of viscous elements into elastic ones, quantified by the exponential decay of *P* (*t*) (Figure 3j). To elucidate the origin of this exponential decay, we use a stochastic model to describe the state transition of elements. Our theory predicts two distinct aging regimes. If temperature is high (i.e., *α >* 1), *P* (*t*) decays exponentially in time; otherwise, *P* (*t*) decays as a power-law function of time with a long-time exponent between 1 and 2.

Our work opens several avenues for future investigation. First, the generality of our model could be tested against other types of condensate systems. Second, molecular dynamic simulations could be employed to establish a direct link between specific molecular features (e.g., sticker density, chain flexibility, interaction valence) and the phenomenological parameters *P* and *f*. In conclusion, our work provides a quantitative, predictive theoretical framework that bridges mesoscopic dynamics with macroscopic rheology, offering a deeper understanding of the physical principles governing the aging behavior of biomolecular condensates.

## Acknowledgments

We thank Huan-Xiang Zhou and Bohan Lyu for helpful discussion related to this work. The research was funded by the National Key Research and Development Program of China (2024YFA0919600), the National Natural Science Foundation of China (Grant No. 12474190) and Peking-Tsinghua Center for Life Sciences grants.

## Notes

### Competing Interest Statement

The authors have declared no competing interest.

