## Supplemental Material for "Mesoscopic viscous-to-elastic transition underlies aging of biomolecular condensates"

### A. Derivation of the Viscoelastic Model

We derive the constitutive equations for the proposed viscoelastic model. Consider a step strain, $\gamma_0$, applied to the system at $t = 0$. At the moment of application ($t = 0^+$), the initial conditions are that the strain on the series elements is $\gamma_s(0^+) = \gamma_0/f$ and on the parallel elements is $\gamma_p(0^+) = 0$. Consequently, the stress $\sigma(0^+) = \gamma_0 G_0/(f(1-P))$. Applying the Laplace transform to the constitutive equations [Eqs. (1) and (2) in the main text] and using the condition $\gamma(t) = \gamma_0$ for $t \geq 0$, we obtain the constitutive relationship in the frequency domain:

$$\begin{pmatrix} (1-f)\,s & f\left(\frac{1-P}{G_0}s + \frac{P}{\eta_0}\right) \\ -\left[P\eta_0 s + (1-P)\,G_0\right] & 1 \end{pmatrix} \begin{pmatrix} \gamma_p(s) \\ \sigma(s) \end{pmatrix} = \begin{pmatrix} \gamma_0 \\ 0 \end{pmatrix}, \tag{S1}$$

where $s$ is the complex frequency, $\sigma(s)$ is the Laplace-transformed stress, and $\gamma_p(s)$ is the Laplace-transformed strain of the parallel section. Solving Eq. (S1) leads to

$$\sigma(s) = G_0\gamma_0 \frac{1}{f[\frac{P}{\tau_0} + (1-P)s] + \frac{(1-f)s}{Ps\tau_0 + 1 - P}}. \tag{S2}$$

A straightforward calculation shows that the above equation can be written as

$$\frac{\sigma(s)}{\gamma_0} = \frac{G_1}{s + \frac{1}{\tau_1}} + \frac{G_2}{s + \frac{1}{\tau_2}}, \tag{S3}$$

with $G_1$, $G_2$, $\tau_1$ and $\tau_2$ defined in Eqs. (3a) and (3b) of the main text. Eq. (S3) corresponds to a double-exponential decay in real time, i.e., two Maxwell fluids connected in parallel.

The characteristic relaxation time, $\tau_c$, is defined as the ratio of the viscosity to the instantaneous modulus:

$$\tau_c = \frac{\eta}{G(t=0)} = \frac{G_1}{G_1 + G_2}\tau_1 + \frac{G_2}{G_1 + G_2}\tau_2 = \frac{1-P}{P}\frac{\eta_0}{G_0}. \tag{S4}$$

This relation predicts that in the limit $P \to 0$, the relaxation time diverges as $\tau_c \sim 1/P$. We note that the system's relaxation time $\tau_c$ is the weighted average of the fast $\tau_1$ and slow $\tau_2$ modes. As $P \to 0$, $(G_2\tau_2/G_1\tau_1) \sim 1/[f(1-f)P^2]$. Consequently, $\tau_2$ dominates $\tau_c$.

We note that $\tau_1$ and $\tau_2$ intersect at $P = 0.5$ when $f = 1$. $G_1$ and $G_2$ intersect at $P_c = (2f-1)/(2f)$ for $f \geq 1/2$. In the case of $f = 1$, the viscoelastic model reduces to a single Maxwell fluid, as $\tau_1$, $\tau_2$, $G_1$, and $G_2$ become piecewise-defined functions (Figure 1c, d of the main text):

$$\tau_1 = \begin{cases} \dfrac{1-P}{P}\tau_0, & P > 0.5, \\ \dfrac{P}{1-P}\tau_0, & P < 0.5, \end{cases} \qquad \tau_2 = \begin{cases} \dfrac{P}{1-P}\tau_0, & P > 0.5, \\ \dfrac{1-P}{P}\tau_0, & P < 0.5, \end{cases} \tag{S5}$$

$$G_1 = \begin{cases} 0, & P < 0.5, \\ \dfrac{1}{1-P}G_0, & P > 0.5, \end{cases} \qquad G_2 = \begin{cases} \dfrac{1}{1-P}G_0, & P < 0.5, \\ 0, & P > 0.5. \end{cases} \tag{S6}$$

#### B. Derivation of the Aging Dynamics

We apply the Laplace transform to Eq. (8) of the main text with the initial condition $p_e(E, t = 0) = 0$. In the Laplace space, the conservation condition $P(t) + \int_0^\infty p_e(E, t)\mathrm{d}E = 1$ becomes $P(s) + \int_0^\infty p_e(E, s)\mathrm{d}E = 1/s$, and we find that

$$sp_e(E, s) = -p_e(E, s)\Gamma_0 e^{-\beta E} + \left[\Gamma_0\beta_0 e^{-\beta_0 E} + \Gamma_1\delta(E - E^*)\right]\left(\frac{1}{s} - \int_0^\infty p_e(E', s)\mathrm{d}E'\right). \tag{S7}$$

Integrating Eq. (S7) over energy barriers gives the total elastic probability $P_e(s) = \int_0^\infty p_e(E, s)\mathrm{d}E$,

$$P_e(s) = \frac{1}{s}\frac{C(s)}{1 + C(s)}, \tag{S8}$$

where $C(s) = \int_0^\infty (\Gamma_0\beta_0 e^{-\beta_0 E} + \Gamma_1\delta(E - E^*))/(s + \Gamma_0 e^{-\beta E})\mathrm{d}E$. The viscous probability $P(s)$ is then found from the normalization condition $P(s) + P_e(s) = 1/s$, giving

$$P(s) = \frac{1}{s(1 + C(s))}. \tag{S9}$$

In the following, we study the asymptotic behavior of $P(s)$ in the limit $s \ll 1$.

##### *I. Case $\alpha > 1$*

For $\alpha \equiv \beta_0/\beta > 1$, $\int_0^\infty \Gamma_0\beta_0 e^{-\beta_0 E}/(s + \Gamma_0 e^{-\beta E})\mathrm{d}E \to \alpha/(\alpha - 1)$ in the limit of $s \ll 1$; therefore, we obtain

$$P(s) \approx \frac{1}{s(\frac{2\alpha-1}{\alpha-1} + \frac{\Gamma_1}{s+\Gamma_0 e^{-\beta E^*}})} \approx \frac{1}{s(\frac{2\alpha-1}{\alpha-1} + \frac{\Gamma_1}{s})}. \tag{S10}$$

Here, we neglect the term $\Gamma_0 \exp(-\beta E^*)$ since $\beta E^* \gg 1$. Eq. (S10) leads to the long-time behavior of $P(t)$ as

$$P(t) \sim e^{-\frac{\alpha-1}{2\alpha-1}\Gamma_1 t}. \tag{S11}$$

One can also find the plateau value $P = (\alpha - 1)/(2\alpha - 1)$ before reaching the exponential decay regime by taking $\Gamma_1 = 0$ in Eq. (S10).

##### *II. Case $\alpha < 1$*

For $\alpha < 1$, $\int_0^\infty \Gamma_0\beta_0 e^{-\beta_0 E}/(s + \Gamma_0 e^{-\beta E})\mathrm{d}E \to \alpha(s/\Gamma_0)^{\alpha-1}\pi\csc(\alpha\pi)$ in the limit of $s \ll 1$. Thus, $P(s)$ becomes

$$P(s) \approx \frac{1}{s\left(1 + \alpha(s/\Gamma_0)^{\alpha-1}\pi\csc(\alpha\pi) + \frac{\Gamma_1}{s+\Gamma_0 e^{-\beta E^*}}\right)}. \tag{S12}$$

In the limit of $\beta E^* \gg 1$, we find the asymptotic behavior of $P(s)$ as

$$P(s) \to \frac{1}{\Gamma_1}(1 - A\frac{s^\alpha}{\Gamma_1\Gamma_0^{\alpha-1}}), \tag{S13}$$

where $A$ is a dimensionless constant, and the corresponding long-time behavior becomes

$$P(t) \sim \frac{1}{\Gamma_0^{\alpha-1}\Gamma_1^2}\frac{1}{t^{1+\alpha}}. \tag{S14}$$

One can find the early-time scaling by taking $\Gamma_1 = 0$ in Eq. (S12) such that $P(S) \to s^{-\alpha}$, leading to $P(t) \sim (\Gamma_0 t)^{\alpha-1}$.

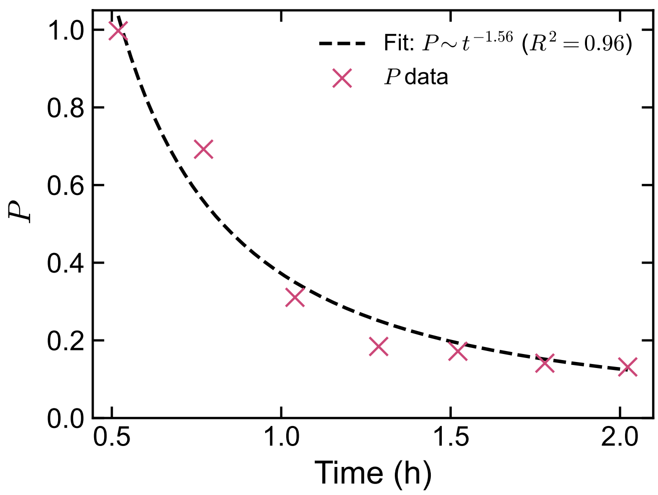


FIG. S1. Power-law fitting of the parameter $P$ as a function of the waiting time. The data can be well fitted by a power-law function with exponent $1 + \alpha \approx 1.56$, consistent with the theoretical prediction for $\alpha < 1$.
